# Yeast *de novo* genes preferentially emerge from divergently transcribed, GC-rich intergenic regions

**DOI:** 10.1101/119768

**Authors:** Nikolaos Vakirlis N, Alex S Hebert, Dana A Opulente, Guillaume Achaz, Chris Todd Hittinger, Gilles Fischer, Josh J Coon, Ingrid Lafontaine

## Abstract

New genes, with novel protein functions, can evolve “from scratch” out of intergenic sequences. These *de novo* genes can integrate the cell’s genetic network and drive important phenotypic innovations. Therefore, identifying *de novo* genes and understanding how the transition from noncoding to coding occurs are key problems in evolutionary biology. However, identifying *de novo* genes is a difficult task, hampered by the presence of remote homologs, fast evolving sequences and erroneously annotated protein coding genes. To overcome these limitations, we developed a procedure that handles the usual pitfalls in *de novo* gene identification and predicted the emergence of 703 *de novo* genes in 15 yeast species from two genera whose phylogeny spans at least 100 million years of evolution. We established that *de novo* gene origination is a widespread phenomenon in yeasts, only a few being ultimately maintained by selection. We validated 82 candidates, by providing new translation evidence for 25 of them through mass spectrometry experiments. We also unambiguously identified the mutations that enabled the transition from non-coding to coding for 30 *Saccharomyces de novo* genes. We found that *de novo* genes preferentially emerge next to divergent promoters in GC-rich intergenic regions where the probability of finding a fortuitous and transcribed ORF is the highest. We found a more than 3-fold enrichment of *de novo* genes at recombination hot spots, which are GC-rich and nucleosome-free regions, suggesting that meiotic recombination would be a major driving force of *de novo* gene emergence in yeasts.

## Introduction

How new genes originate is a fundamental question in evolution. The mechanism of gene acquisition by *de novo* emergence from previously non-coding sequences, has long been considered as highly improbable (Jacob 1977; Kaessmann 2010). New genes were assumed to appear mostly from previously existing coding sequences, through duplication and divergence (Ohno 1970), horizontal transfer (Lerat et al. 2005), or through chimerism, see (Long et al. 2003; Bornberg-Bauer et al. 2010; Kaessmann 2010; Andersson et al. 2015) for reviews. However, for the last decade, a handful of *de novo* genes have been functionally characterized in all eukaryotic lineages, exemplifying their contribution to evolutionary innovations and their integration into central cellular functions (Begun et al. 2006; Levine et al. 2006; Begun et al. 2007; Zhou et al. 2008; Cai et al. 2008; Knowles and McLysaght 2009; Li et al. 2010) and (Wu and Zhang 2013; McLysaght and Guerzoni 2015).

By definition, a *de novo* gene that emerged in a given genome is taxonomically restricted to that single species or, if it originated before speciation, to a group of closely related species. However, Taxonomically Restricted Genes (TRG) also include highly diverged homologs, horizontally acquired genes from yet unsampled species and dubious Open Reading Frames (ORF) erroneously annotated as protein coding genes (Khalturin et al. 2009; Tautz and Domazet-Lošo 2011). Conservative approaches used in the first case studies excluded all genes that had homologs, even in closely related species, and restricted the *de novo* candidates to the genes for which enabling mutations could be retraced from the ancestral non-coding sequence (Begun et al. 2006; Levine et al. 2006; Begun et al. 2007; Zhou et al. 2008; Cai et al. 2008; Knowles and McLysaght 2009; Li et al. 2010). By contrast, large-scale approaches either considered all TRG as *de novo* genes (Carvunis et al. 2011, Neme and Tautz, 2013, Abrusan 2013), or classified TRG based on their probable origin (Donoghue et al. 2011). The issue of false positive TRG detection is therefore problematic, resulting in gene age underestimation, a matter still being debated (Moyers and Zhang 2014 Oct 13; McLysaght and Hurst 2016; Moyers and Zhang 2016; Domazet-Lošo et al. 2017; Moyers and Zhang 2017). Therefore, the quantitative importance of *de novo* gene emergence and their evolutionary dynamics remain poorly understood.

Another open question in the field of gene origination is how DNA sequences undergo transition from noncoding to coding. In order for that to happen, the non-coding region needs to gain two properties: first, become an ORF and then being transcribed, or the other way round. The resulting mRNA molecule must be translated and the protein must enter into the cellular metabolism (Bornberg-Bauer et al. 2015; McLysaght and Guerzoni 2015; Schlötterer 2015). The RNA-first model, in which the formation of an ORF occurs in a region that is already transcriptionally active, is supported by previous reports on *de novo* genes (Cai et al. 2008; Zhou et al. 2008) and by both pervasive transcription (Nagalakshmi et al. 2008; Djebali et al. 2012; Ruiz-Orera et al. 2014; Neme and Tautz 2016) and pervasive translation (Wilson and Masel 2011; Ingolia et al. 2014; Ji et al. 2015). Furthermore, the onset of transcription can be favored by pre-existing regulatory sequences (Knowles and McLysaght 2009; Siepel 2009) and notably by divergent transcription from bidirectional promoters (Core et al. 2008; Neil et al. 2009) which were shown to facilitate new gene origination (Gotea et al. 2013; Neme and Tautz 2013; Wu and Sharp 2013). However, this is not always necessary, as potentially coding *de novo* transcripts are not systematically enriched in bidirectional promoters in humans (Ruiz-Orera et al. 2015), nor in *Drosophila*, where it has been shown that regulatory regions can also emerge *de novo*, along with new genes (Zhao et al. 2014).

Whether every noncoding sequence in a genome has the potential to evolve into a protein-coding gene is another crucial point of interest. In the continuum hypothesis, *de novo* gene birth is a gradual maturation process, from a pool of random non-coding sequences to fully mature genes (Carvunis et al. 2011), while in the preadaptation hypothesis, *de novo* genes emerge by a non-gradual process, only within pre-adapted genomic regions, that harbor gene-like characteristics (Masel 2006; Wilson et al. 2017).

In addition, independently of the mode of transition (RNA-first or ORF-first) or from its origination route (continuum or preadaptation hypotheses), a *de novo* gene first emerges at a low frequency in a population and subsequently can either disappear or eventually reach fixation. Few population analyses showed that the life cycle of *de novo* genes would be relatively short and may depend on lineage-specific functional requirements (Palmieri et al. 2014; Zhao et al. 2014; Li et al. 2016).

Here, we developed a multi-level systematic approach which addresses all the above issues and strikes a balance between previously published, broader proto-gene surveys (Carvunis et al. 2012) and stricter, but more limited approaches such as the ones applied in human (Knowles and McLysaght 2009). We used this approach to search for *de novo* genes in two yeast genera comprising a total of 15 species, which span at least 100 million years of evolution (Berbee and Taylor 2006). Based on our results, we propose a plausible mechanistic model of *de novo* gene emergence in yeasts, in which meiotic recombination plays a crucial role.

## Results

### A comprehensive methodology for a reliable genus-wide identification of de novo gene candidates

We developed a novel approach to reliably identify *de novo* gene candidates at the genus level and applied it to two yeast genera with high quality genome assemblies: the genus *Lachancea* (Kellis et al. 2004; Souciet et al. 2009; Vakirlis et al. 2016)□ and the well characterized genus *Saccharomyces* [(Scannell et al. 2011), Fig. S1 and Table S1].

We first identified 1837 TRG, *i.e.* with no detectable homologs outside of each of the two genera after clustering all annotated CDS into singletons and homologous families, and performing an exhaustive similarity search with both single sequence-based and profile-based tools against several public databases. Then we inferred their branch of origin along the genus phylogeny by phylostratigraphy [see Methods, (Domazet-Lošo et al. 2007; Domazet-Lošo et al. 2017)]. Second, we eliminated 55 fast-diverging TRG families in *Lachancea* (no *Saccharomyces* TRG families were removed at this step), whose ages are most likely underestimated according to the expected number of false positive predictions given an evolutionary distance between homologs of simulated protein families (see Fig. S2, Fig. S3 and Methods). Third, we filtered out 1028 TRG that are more likely to be spurious ORFs than true protein-coding genes. To this end, we developed a logistic regression classifier trained on codon usage and sequence-based properties of known noncoding sequences. The classifier assigns a statistical Coding Score (CS) to each TRG. We defined a genus-specific CS threshold above which we can expect only 5% of non-coding sequences erroneously classified as coding and removed 701 and 327 TRG with lower CS than the thresholds in *Saccharomyces* and in *Lachancea*, respectively (see Fig. 1A, Fig. S4, Table S2 and Methods).

**Figure 1.**
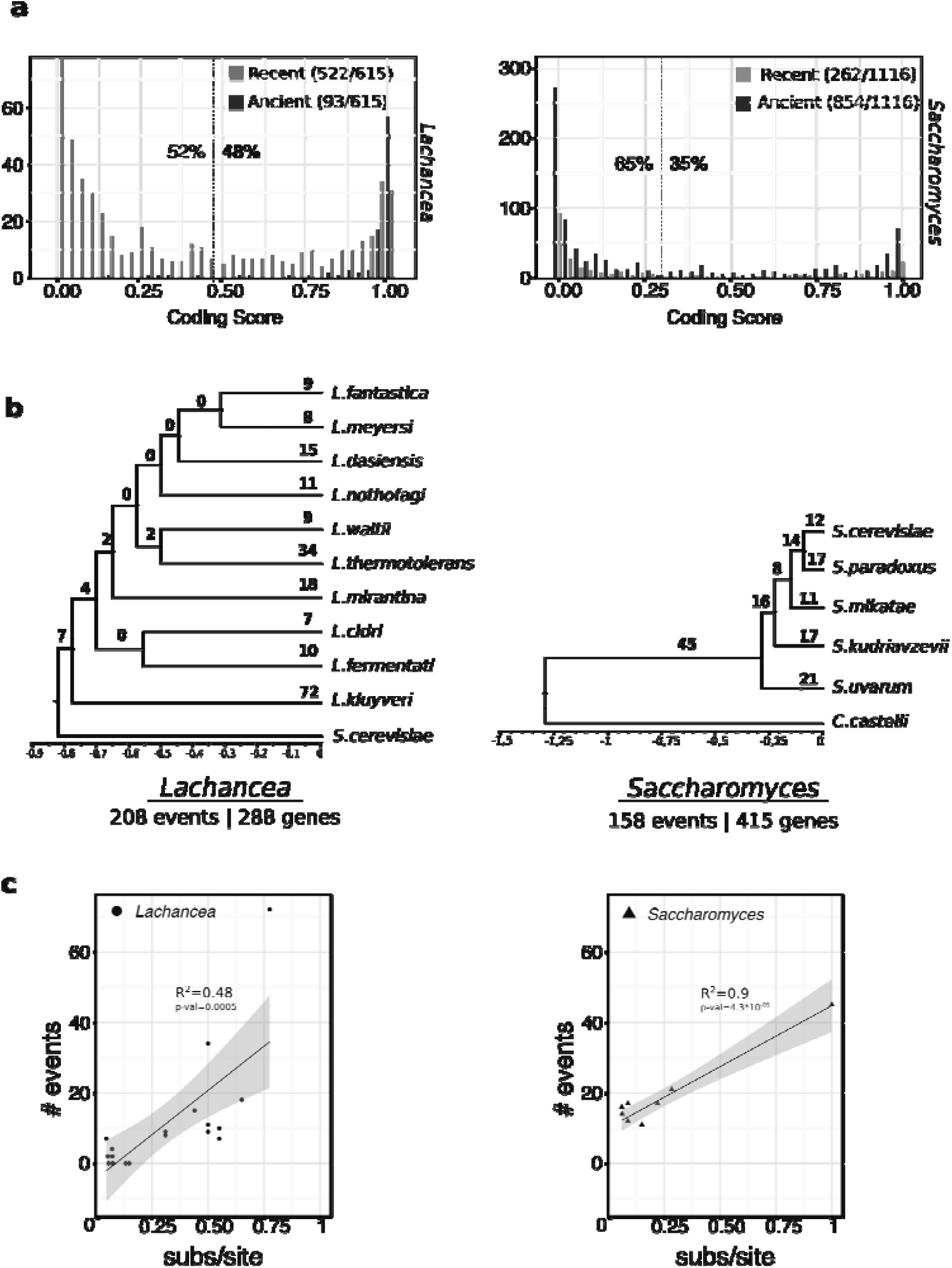
Results of *de novo* gene identification in 2 model yeast genera. (**A**) Distributions of Coding Scores (CS) of TRGs in the 2 genera. Dashed lines represent thresholds (0.47 in *Lachancea,* 0.3 in *Saccharomyces*) that limit false positives to 5% based on our validation procedure (Figure S1 and Methods). Black bars: ancient TRG, grey bars: recent TRG (**B**) *De novo* gene origination along the phylogenies of the 2 genera. Branch lengths correspond to molecular clock estimations of relative species divergence (relative number of substitutions per site) within each genus. Thus, the bottom scale bar expresses species relative number of substitutions per site to the origin of the genus. Recent and ancient events are shown in red and green, respectively. (**C**) Numbers of *de novo* creation events as a function of the relative number of substitutions per site to the origine of the genus, as shown in **B.**

### A robust set of yeast de novo gene candidates

Based on the above methodology, we selected 701 *de novo* gene candidates, which were derived from an estimated total of 366 events of *de novo* gene creation that took place during the evolution of the two genera (Fig. 1B, Table S3). For further analysis, we added 2 previously described *de novo* genes in *S. cerevisiae,* BSC4 (Cai et al. 2008) whose CS was below the genus threshold, and MDF1 (Li et al. 2010) which was not annotated in the version of the *S. cerevisiae* genome that we used (Scannell et al. 2011). We considered the *de novo* candidates as being “recent” when they were restricted to one species, *i.e.* when they emerged along a terminal branch of the phylogenetic tree or “ancient” when they emerged along an internal branch of the tree (Fig. 1A and 1B). Taken together, the 288 and 415 *de novo* gene candidates respectively account for 0.45% and 0.9% of the gene repertoire in *Lachancea* and *Saccharomyces*.

We found that, in both genera, branch lengths correlated to the number of *de novo* origination events (Fig. 1C) suggesting that *de novo* emergence occurs at a coordinated pace with non-synonymous mutations. However, these results are best viewed qualitatively given the limited number of data points: the slopes of the fitted regression lines are unlikely to represent the true emergence rates. There is a smaller estimated time for the divergence between the *Nakaseomyces/Candida* and the *Saccharomyces* genus -from 57 to 87 Mya-than between the *Kluyveromyces/Eremothecium* and the *Lachancea* genus - from 84 to 126 Mya - (Kensche et al. 2008; Doyon et al. 2012; Beimforde et al. 2014; Marcet-Houben and Gabaldón 2015). However, the average number of events per lineage since the divergence of the *Saccharomyces* is significantly greater (83.8) that the one since the divergence of the *Lachancea* (31.7), (p=0.0058, Wilcoxon test). Similarly, the average number of origination events per substitution per site for each branch (*i.e*. branch length of the trees in Figure 1B) is 133.8 in *Saccharomyces* and 32.7 in *Lachancea* (p=0.002 Wilcoxon test). Both exogenous (different environmental or selective pressures) and intrinsic (differences in genome dynamics) factors could account for these variations.

### Experimental validation of de novo proteins

We provide experimental evidence of translation for 25 *de novo* genes in *Lachancea* by performing tandem mass spectrometry (MS/MS) analysis at the whole proteome level in rich growth medium conditions (Table S2, Table S3 and Methods). Prior global proteomic experiments in *S. cerevisiae* validated 60 out of the 105 *de novo* gene candidates in that species (Table S4). Altogether, experimental evidence of translation validates 85 (12%) of our candidates, which we will refer to as validated *de novo* genes hereafter. Crucially, the CS of the validated *de novo* genes is very high (median at 0.95). Conversely, in the *Lachancea* species, we found that none of the TRG eliminated as spurious (based on their low CS) was detected by MS. Among the 302 CDS that we classified as spurious TRG in *S. cerevisiae*, only 13 show evidence of translation, thus likely corresponding to false negatives that were misclassified by our logistic regression classifier. The median CS of the 85 validated *de novo* genes plus the 13 validated spurious TRG is 0.8, suggesting that our CS is a good indicator of protein expression (see Methods).

In both genera, there are less recent than ancient validated *de novo* genes, with only three (12%) in *Lachancea*, and five (20%) in *Saccharomyces*, suggesting that recent *de novo* genes are poorly expressed. Among the validated *de novo* genes in *S. cerevisiae,* four have a known function: *REC104* and *CSM4* are involved in meiotic recombination, *PEX34* is involved in the peroxisome organization, and *HUG1* participates to the response to DNA replication stress (Table S4).

### The transition from non-coding to coding can be inferred for 30 Saccharomyces de novo genes

The most convincing evidence of *de novo* gene birth stems from the unambiguous identification of the mutations that enabled the formation of an open reading frame in a given lineage, when compared to the orthologous non-coding regions closely related genomes. Based on multiple alignments between the *de novo* genes and their orthologous DNA sequences in closely related genomes as in (Knowles and McLysaght 2009) (see Methods), we identified one or several ancestral nucleotide(s) that once mutated, gave rise to the ORF for 30 *de novo* genes in *Saccharomyces* (Fig. 2). Among these 30 *de novo* genes that we hereafter label “reliable”, 27 belong to the “recent” group and show higher similarity to their orthologous intergenic regions compared to the genomic average. Therefore, they are probably some of the most recently emerged ones.

**Figure 2.**
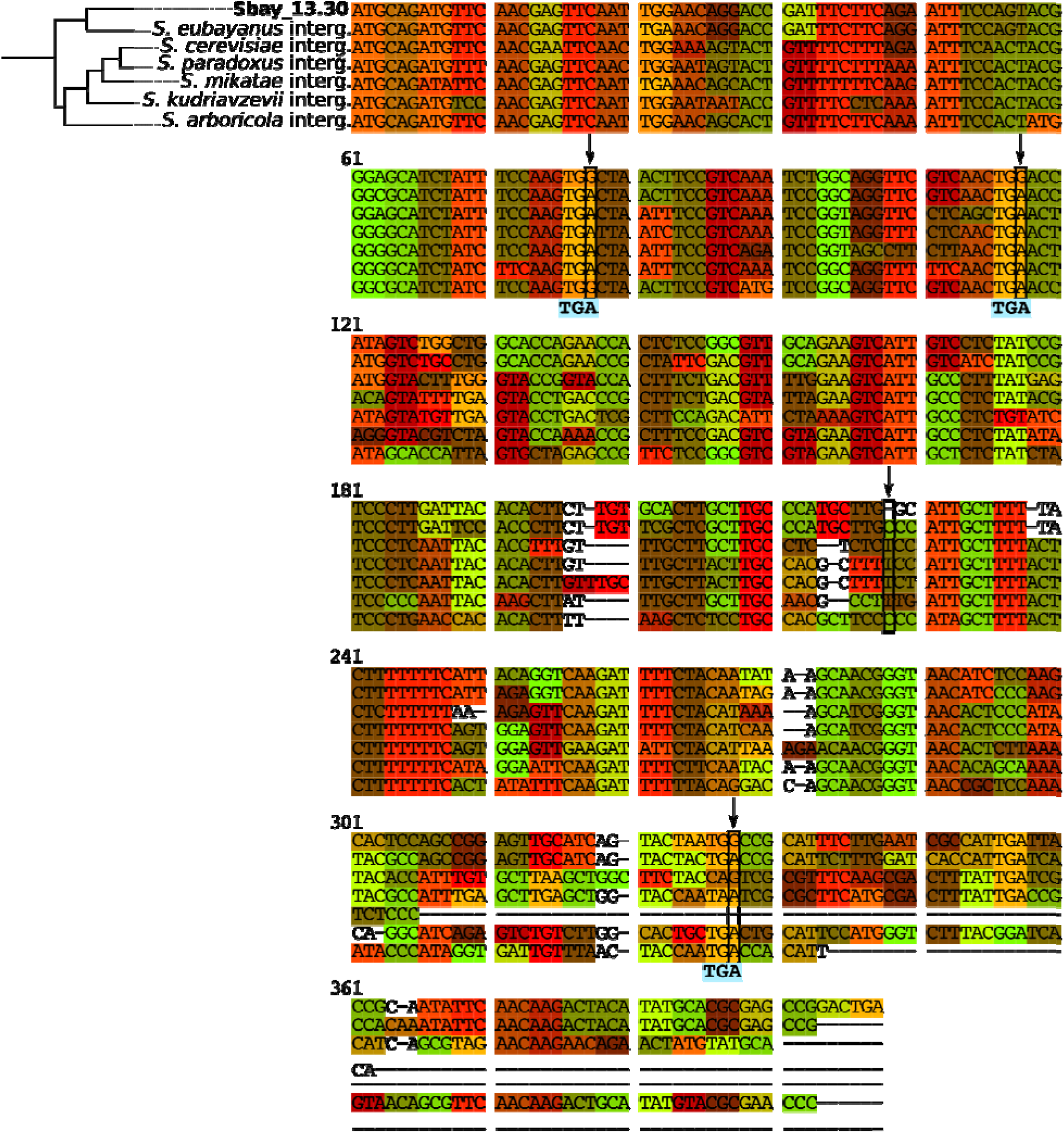
Alignment of the *de novo* gene Sbay_13.30 in *S. uvarum* and its orthologous intergenic sequences in all other *Saccharomyces* genomes. Four enabling mutations that occurred along the *S. uvarum* branch are indicated with an arrow. Ancestral states for the critical positions are shown under the alignment (positions were the same at the root of the *Saccharomyces* and the common ancestor of *S. uvarum* and *S. eubayanus*). At least 3 stop codons were removed by base substitution and a frameshift occurred due to the deletion of one base leading to the formation of the ORF in *S. uvarum*.

No such mutational scenario could be retrieved in the *Lachancea species*, because their genomes are too divergent, with orthologous intergenic regions that no longer share significant similarity.

### De novo genes have unique sequence characteristics as compared to conserved genes

The *de novo* candidates share a number of structural properties that differentiate them from the genes conserved outside the two genera. They are significantly shorter, have a lower codon adaptation index and a higher aggregation propensity compared to conserved genes (Fig. S5). Their biosynthetic cost is also lower than those of non-coding sequences, in agreement with an intermediate stage from a non-coding to a coding state (Fig. S5). When recent, *de novo* genes are not enriched in intrinsically disordered regions compared to conserved genes. The low propensity of recent genes to disorder was previously reported in *S. cerevisiae* (Carvunis et al. 2012). When ancient, but in *Lachancea* only, *de novo* genes have a higher proportion of predicted disorder than conserved genes (Fig. 3), suggesting contrasted evolutionary pressures (see Discussion Section).

**Figure 3.**
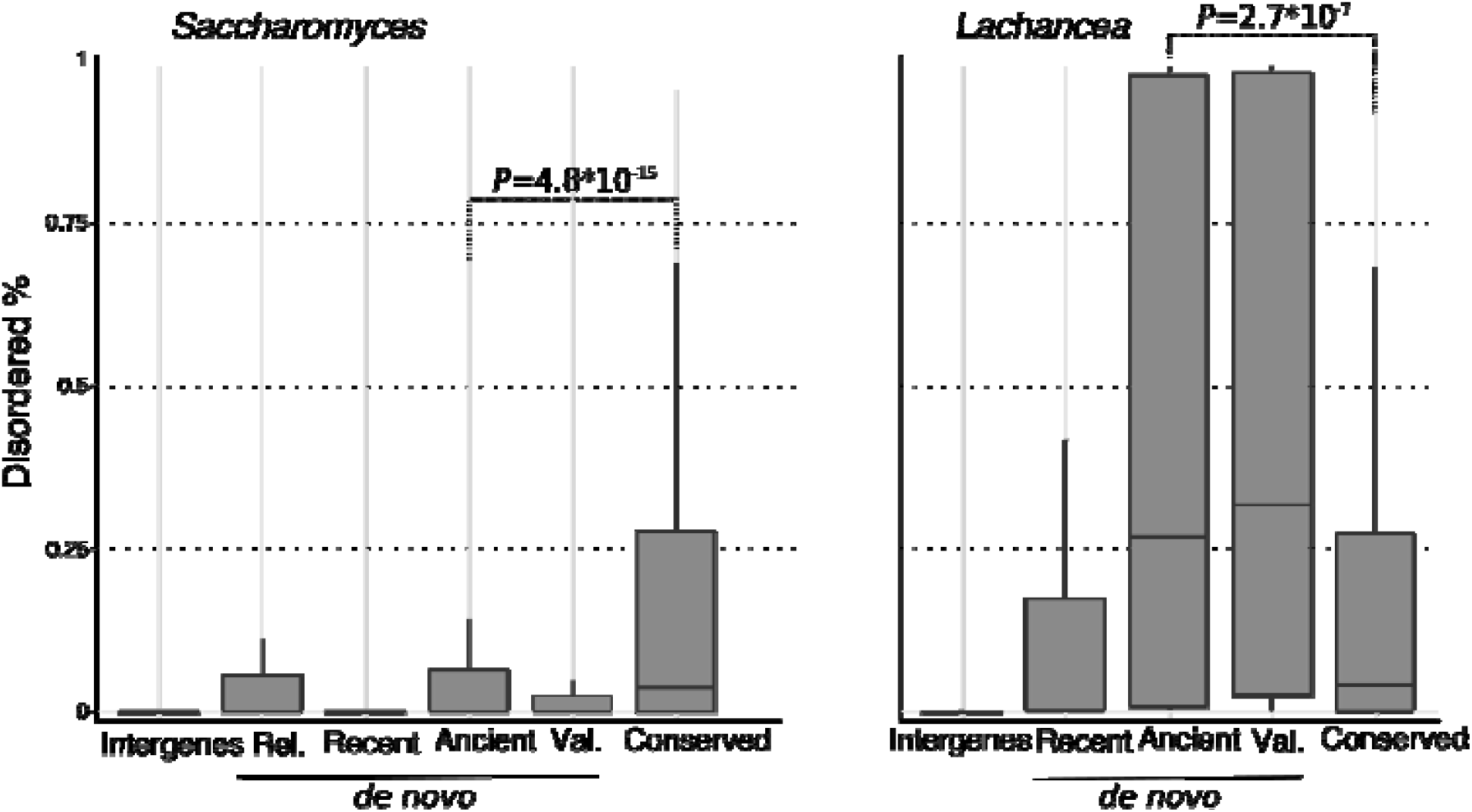
Distributions of percentages of residues in disordered regions for various sequence classes in the 2 yeast genera. Rel.: reliable *de novo* genes for which the ancestral sequence is inferred as non-coding. Val.: validated *de novo* genes with experimental translation evidence.

### De novo genes preferentially emerge next to divergent promoters in GC-rich intergenic regions

We found that *de novo* genes are significantly enriched in opposing orientation with respect to their direct 5’ neighboring gene (Fig. 4A). Similar enrichment was already observed for mouse-specific genes (Neme and Tautz 2013). This suggests that *de novo* genes would benefit from the divergent transcription initiated from bidirectional promoters. By contrast, tandemly duplicated genes are significantly enriched in co-orientation with respect to their 5’neighbour (69% and 74% in *Saccharomyces* and *Lachancea*, respectively (not shown)). Therefore, the bias towards opposing orientations strongly suggests that the *de novo* gene candidates do not actually correspond to tandemly duplicated genes that would have diverged beyond recognition. In addition, the bias towards divergent orientation is the strongest for the reliable *de novo* genes which correspond to the most recently emerged genes (see above), suggesting that divergent transcription from bidirectional promoters is critical in the early stages of origination.

**Figure 4.**
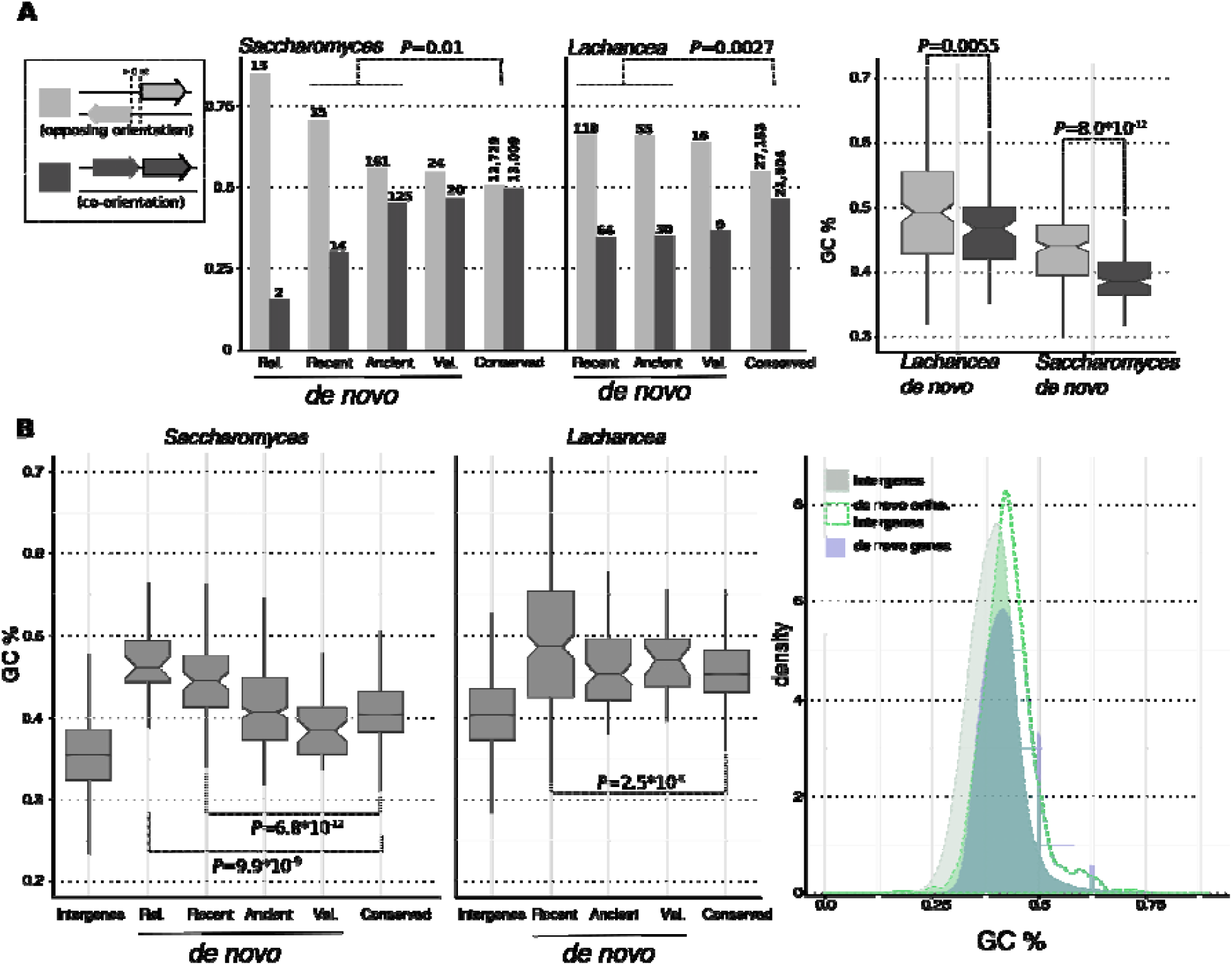
*De novo* genes are enriched at divergent promoters in GC-rich regions. (**A**) Left and middle: Distributions of the transcriptional orientations of various gene classes relative to their 5’ neighbours (see text). Only genes with a non-null 5’ intergenic spacer (> 0 nt) are considered. Right: GC% distributions of *de novo* genes in opposing and co-orientation configurations in the 2 genera. Grey: opposing orientation, black: co-orientation. (**B**) Distributions of Guanine-Cytosine percentage (GC%) in various sequence classes. Notches represent the limits of statistical significance. De novo ortho. intergenes: intergenic regions orthologous to *de novo* genes. Rel: reliable, Val.: validated.

Recent and reliable *de novo* genes in *Saccharomyces* and recent ones in *Lachancea* have significantly higher GC content than conserved genes, which are themselves more GC-rich than intergenic regions (Fig. 4B, Fig. S6B and S6C). Moreover, *de novo* genes in opposing orientation with respect to their 5’ gene neighbor are also more GC-rich than co-oriented ones (Fig. 4A right). Finally, we found that the orthologous noncoding regions of *de novo* genes in sister genomes have a GC content significantly higher than that of the other intergenic regions (Fig. 4B right, Figure S6A, Table S5). Therefore we propose that *de novo* genes tend to emerge in particularly high GC-rich regions, where the frequency of AT-rich stop codons is the lowest and the probability of finding a fortuitous and transcribed ORF is therefore the highest (Figure S6A).

### De novo genes are significantly enriched at recombination hotspots

In multiple eukaryotic taxa, including yeasts and humans, heteroduplexes formed during meiotic recombination are repaired by gene conversion biased towards GC-alleles, thus increasing the GC-content of recombination hotspots (RHS) (Lamb 1984; Jeffreys and Neumann 2002; Mancera et al. 2008; Duret and Galtier 2009). Furthermore, it provides a nucleosome-free region (Berchowitz et al. 2009; Pan et al. 2011) that promotes transcriptional activity. It follows then that RHS could be favorable locations for the emergence of *de novo* genes in yeasts. We tested enrichment of *de novo* genes overlapping with RHS in *S. cerevisiae*, *S. mikatae* and *S. kudriavzevii*, the species for which recombination maps are exploitable for this study [(Lam and Keeney 2015a), see Methods] (Fig 5A). The enrichment was tested against i) *de novo* genes overlapping with a set of randomly shuffled hotspot-equivalent regions and ii) a set of conserved genes (with the same GC content, length and chromosome distributions as *de novo* genes) overlapping with the real RHS (P-value<0.001 calculated from 1000 simulations for all tests, except for *S. kudriavzevii* in the sampled-conserved test, P-value = 0.012). More than a third of *de novo* genes overlap with RHS (44%, 42% and 39% *S. cerevisiae*, *S. mikatae* and *S. kudriavzevii*, respectively), which represents a more than 3-fold enrichment (Fig. 5A). The *de novo* genes associated with RHS include 3 validated *de novo* genes in *S. kudriavzevii* and 3 in *S. mikatae*. The length coverage of the *de novo* genes by RHS is on average 65% (204 nt), 66% (192 nt) and 42% (178 nt) in *S. cerevisiae*, *S. mikatae* and *S. kudriavzevii*, respectively. Such a strong association suggests that gene conversion biased towards GC-alleles during meiotic recombination would be a major driving force of *de novo* gene emergence in yeasts.

**Figure 5.**
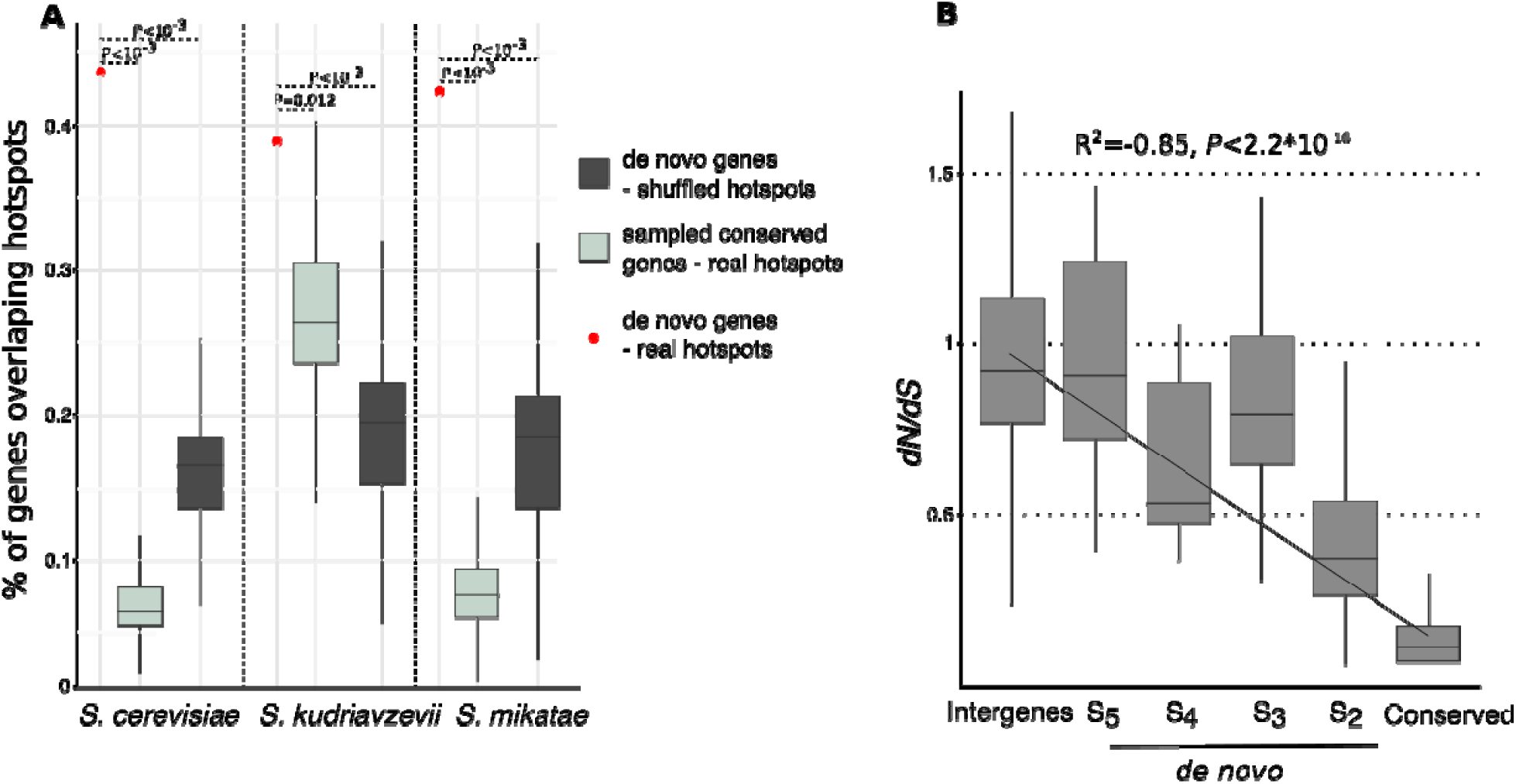
*De novo* genes are enriched at recombination hotspots and are under increasing purifying selection with age. (A) Proportion of *de novo* genes overlapping recombination hotspots as identified in (Lam and Keeney 2015) (outliers are not shown). The 2 null models consist in i) randomly shuffling the hotspots on each chromosome and ii) sampling a set of conserved genes with the same GC composition and chromosome distribution as *de novo* genes. Both models were repeated 1000 times. Red dots: real *de novo* genes, dark (B) Distribution of pairwise dN/dS value for various sequence classes in *Saccharomyces*. S2 to S5 refer to the branches of emergence of *de novo* genes (see Fig. S3).

### The strength of purifying selection acting on de novo genes increases with age

In *Lachancea* species for which several strains are available, the inferred pN/pS (non-synonymous to synonymous polymorphism rates) ratio is on average significantly lower for *de novo* candidates (Fig. S7). The same trend is observed when comparing species instead of strains (Fig. S8). However, the correlation between pN/pS and CS values is lower than for dN/dS and CS values (Fig. S7 and S8) suggesting that most of the species-specific *de novo* candidates (the most recent ones) are under weak purifying selection, as already observed in yeasts, primates and flies (Cai and Petrov 2010; Carvunis et al. 2012; Palmieri et al. 2014; Zhao et al. 2014; Li et al. 2016).

In *Saccharomyces,* the dN/dS ratio (non-synonymous to synonymous substitution rates) of the most recent *de novo* genes is close to 1 and gradually decreases down to the level of the conserved genes for the most ancient ones (Fig. 5B). This indicates that the strength of purifying selection increases with gene age (data are insufficient in *Lachancea*, see Methods).

## Discussion

To our knowledge, our work represents a unique attempt to tackle the three main issues affecting *de novo* gene identification, namely exhaustive similarity searches, estimation of sequence divergence beyond recognition and erroneous gene annotations. In addition, we inferred the mutations enabling the transition from non-coding to coding sequences for 30 *Saccharomyces de novo* genes and showed that these reliable candidates share the same properties as the rest of the dataset. We also found that *de novo* gene candidates were on average smaller than conserved genes and smaller than all documented horizontally transferred genes (Table S5) (Rolland et al. 2009; Marcet-Houben and Gabaldón 2010; Vakirlis et al. 2016). In addition, horizontally transferred genes are predominantly found in co-orientation with respect to their 5’ gene neighbor while candidate *de novo* genes are predominantly found in opposing orientation. Altogether these results show that the set of *de novo* gene candidates identified in this study is mostly devoid of highly divergent homologs, genes horizontally acquired from unknown genomes or neo-functionalized duplicates.

The role of *de novo* emergence as a potent gene birth mechanism has been much debated during the past decade. In this study, we identified a significant number of 701 *de novo* genes candidates (30 of which have unambiguously emerged from ancestral, non-coding sequence) across an unprecedented number of 15 yeasts genomes. Although *de novo* origination occurs at a slow pace, it is sufficiently widespread for *de novo* genes to be present in all genomes studied. In total, the 85 validated *de novo* genes, which have translation evidence, represent 0.1% of the proteome in yeasts, a much higher proportion than what was estimated in other lineages, with 0.01% in *Drosophila*, 0.03% in primates, and 0.06% in the sole *Plasmodium vivax* genome (Chen et al. 2010; Yang and Huang 2011; Guerzoni and McLysaght 2016). On the contrary, there is a significantly higher proportion (2.8%) of validated *de novo* genes specific to the *Arabidopsis thaliana* genome (Li et al. 2016), revealing contrasting dynamics in different eukaryotic lineages. It is also possible that we underestimated the number of validated candidates in yeasts because additional ORFs could actually be expressed in yet untested conditions.

The higher genomic GC content in *Lachancea* – from 41% to 43% - than in *Saccharomyces* – from 38%to40% - could explain the higher proportion of disordered regions of recent *de novo* candidates in *Lachancea* than in *Saccharomyces*, while different evolutionary pressures between the two genera could explain why ancient *de novo* genes, but in *Lachancea* only, have higher proportion of disorder than conserved genes, although they have a similar GC content. These results are in agreement with the recent analysis of Basile and colleagues (Basile et al. 2017) showing that in recent *de novo* genes, the level of disorder is strongly dependent on the genomic GC content and that this dependency decreases during evolution. In another recent article, Wilson and colleagues (Wilson et al. 2017) showed that the correlation between age and disorder observed in 871 *S. cerevisiae de novo* genes disappears when considering only the 35 ancient *de novo* genes presenting the highest probability to encode a functional protein product. When considering only our 60 validated *de novo* genes from *S. cerevisiae*, we observed the same phenomenon.

Finally, our results suggest a reasonable mechanistic model for the early stages of *de novo* evolution in yeasts: *de novo* emergence of ORFs occurs in GC-rich non-coding regions, where the probability of finding a fortuitous ORF is the highest and preferentially where *de novo* ORF can be transcribed from the divergent promoter of its 5’ neighboring gene (Fig 6). Recombination hotspots are good candidate regions for *de novo* emergence because they have a high GC content (Mancera et al. 2008) and they preferentially localize at promoters in yeasts, but also in dogs, birds, and Arabidopsis (Auton et al. 2013; Choi et al. 2013; Lam and Keeney 2015; Singhal et al. 2015). As the stability of an mRNA molecule increases with its GC content (Kudla et al. 2006; Neymotin et al. 2016), the *de novo* GC-rich transcript will be stable, and could thus be efficiently translated. Consequently, whether the protein product will be beneficial l or harmful to the cell, the *de novo* gene will be either fixed or rapidly lost from the population.

**Figure 6.**
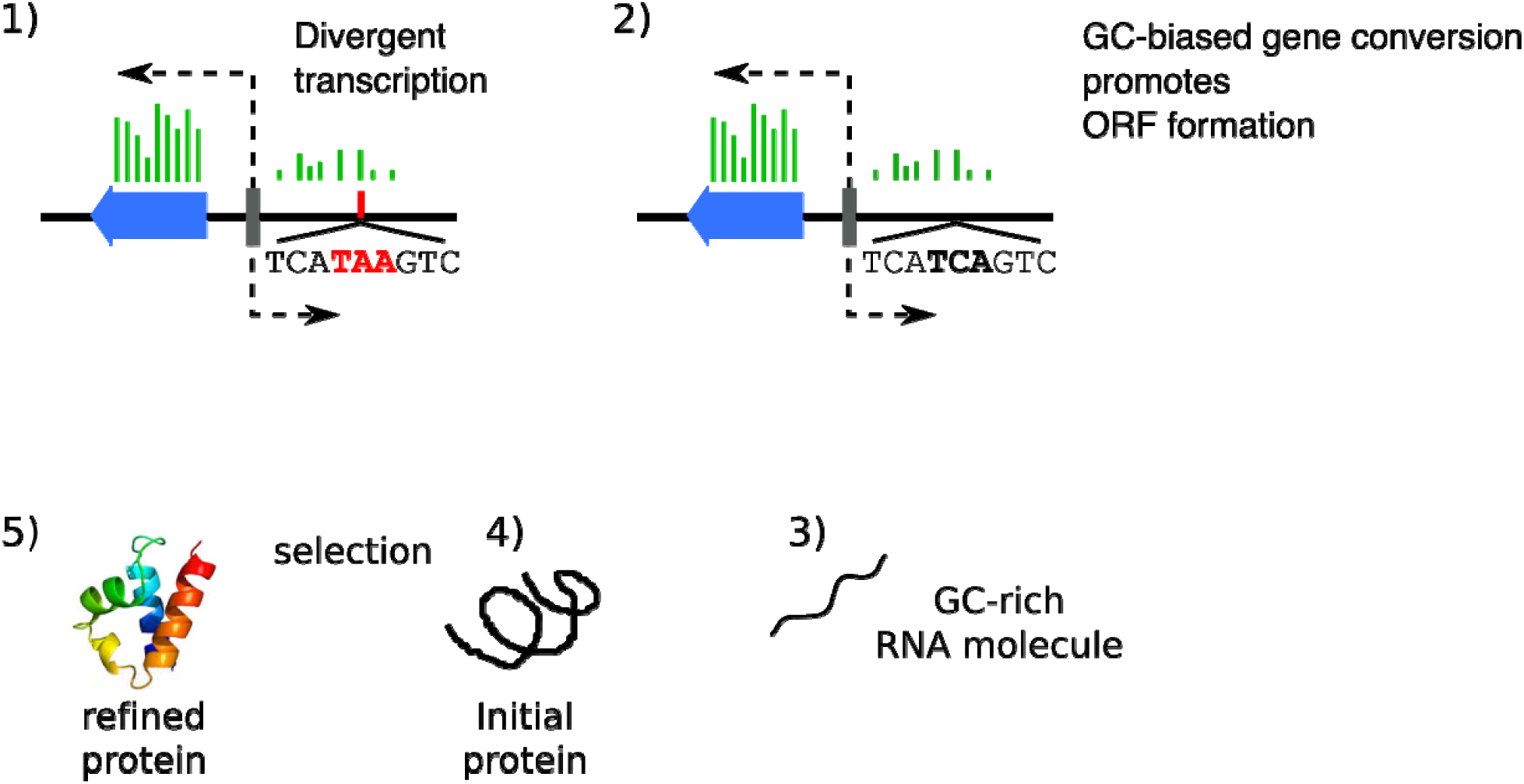
Model of *de novo* gene evolution. Blue arrow: conserved gene. Grey bar: bidirectional promoter. Red bar: stop codon. Green bars: transcription.

We established a new approach to rigorously identify *de novo* gene candidates in a set of closely related species. Interestingly, our study supports a role for meiotic GC biased gene conversion as a driver of gene origination in yeasts. More generally, the role of recombination hostpsots in *de novo* gene emergence in other eukaryotes should prove most interesting to explore in the future.

## Materials and Methods

### Data collection

We investigated *de novo* gene emergence in 10 *Lachancea* and 5 *Saccharomyces* genomes (*L. kluyveri*, *L. fermentati*, *L. cidri*, *L. mirantina*, *L. waltii*, *L. thermotolerans*, *L. dasiensis*, *L. nothofagi*, *"L. fantastica" nomen nudum* and *L. meyersii*, *S. cerevisiae*, *S. paradoxus*, *S. mikatae*, *S. kudriavzevii*, and *S. bayanus var. uvarum*), see Table S1. For the *Saccharomyces*, the genome of *S. arboricola* was not analysed because it contains *ca.* half of the number of annotated genes than the others. It was only used for the reconstruction of the ancestral sequences of *de novo* genes. The genome of *S. eubayanus* was not analysed either because it was not annotated with the same pipeline. It was used for the reconstruction of the ancestral sequences of *de novo* genes and for the simulation of the protein families’ evolution. For outgroup species references, the genomes *Kluyveromyces marxianus*, *K. lactis*, and *K. dobzhanskii* were used for the *Lachancea* and the genomes of *Candida castellii* and *Nakaseomyces bacillisporus* were used for the *Saccharomyces*. The sources for genome sequences and associated annotations are summarized Table S1. Annotated CDS longer than 150 nucleotides were considered.

The high raw coverages of the assembled genomes in the two genera minimized erroneous base calls and make sequencing errors and subsequent erroneous *de novo* assignment very unlikely (N50 values range from 801 to 905 kb for *Saccharomyces* (Scannell et al. 2011) and form 1275 and 2184 kb in *Lachancea*). The combined 454 libraries and Illumina single-reads for the *Lachancea* project (Vakirlis et al. 2016) further allowed the correction of sequencing errors in homopolymer blocks that generated erroneous frameshifts in genes.

### Pipeline for TRG detection

Initially, the protein sequences of all considered species (focal proteome) are compared against each other using BLASTP (Altschul et al. 1990) (version 2.2.28+, with the options *-use_sw_tback - comp_based_stats* and an E-value cut-off of 0.001) then clustered into protein families by TribeMCL (Enright et al. 2002) (version 12-068, I=6.5) based on sequence similarity, as previously reported for the *Lachancea* genomes (Vakirlis et al. 2016). For each family, a multiple alignment of the translated products is generated (see *General procedures* section) and profiles (HMM and PSSM) are built from it. These first steps are also performed for the proteome of the outgroup species.

A similarity search for homologs outside of the focal species is then performed against the NCBI nr database with BLASTP for singletons and with PSI-BLAST version 2.2.28+ for families with their own PSSM profiles. Hits are considered significant if they have an E-value lower than 0.001 for both BLASTP and PSI-BLAST. A family (or singleton) is considered as taxonomically restricted if it has no significant hit in nr. This work was already done for the *Lachancea* in (Vakirlis et al. 2016). TRGs whose coordinates overlapped conserved genes on the same strand were removed.

Next, TRG families are searched against each other using HMM profile-profile comparisons with the HHSUITE programs version 2.0.16 (Söding 2005). HMM profiles were built with *hhmake*, and database searches were performed with *hhsearch*. A hit is considered significant if it has a probability higher than 0.8 and an E-value lower than 1, values previously defined as optimal (Lobb et al. 2015). Families sharing significant similarity are merged. This new set of TRG families is used to search for similarity in 4 databases: an HMM profile database built from the alignments of the genus’ conserved families, the profile database of the outgroup species, the PDB70 profile database, version of 03-10-2016 (Söding et al. 2005), and the PFAM profile database, version 27.0 (Finn et al. 2014). Singleton TRGs were compared by sequence-profile searches using *hmmscan* of the HMMER3 package version 3.1b2 (Mistry et al. 2013) (E-value cut-off 10^−5^) in all the above databases, except PDB70. The final curated TRG families are those for which no significant match is found in any searched database. Finally, the branch of origin of each TRG family is inferred as the branch leading to the most recent common ancestor of the species in which a member of the family is present. The reference species phylogeny is given in Fig. S1.

### Simulations of protein family evolution and removal of false positive TRGs

We simulated the evolution of gene families created before the divergence of the genus along the *Saccharomyces* and *Lachancea* phylogenies. The real orthologous gene families were defined as families of syntenic homologues with only one member per species as in Vakirlis *et al.* (Vakirlis et al. 2016). We defined 3668 such families across the 10 *Lachancea* and their 3 outgroup species, as well as 3946 families across the 6 *Saccharomyces* species and their 2 outgroup species. We followed the simulation protocol used by Moyers *et al.* (Moyers and Zhang 2014; Moyers and Zhang 2016) but we inferred protein evolutionary rates for each individual gene tree (branch lengths representing substitutions per 100 sites), instead of calculating the mean evolutionary rate of a protein by the number of substitutions per site per million years between a couple of yeast species, and we did so using the ROSE program version 1.3 (Stoye et al. 1998) with the PAM matrix. We believe that using a model of protein evolution to detect false positive TRG is reasonable, given that false positive TRG are actually conserved genes that arose before the divergence of the species and thus evolved as protein coding genes for a longer time than *de novo* genes.

We performed simulations under two scenarios. In the first scenario (normal case), the amount of divergence within each simulated protein family mirrors the one within real orthologous families. In the second scenario (worst case), the divergence is 30% higher than the one estimated among the real orthologous families (every simulated branch is 30% longer than its real equivalent), and additionally, for each branch, a random amount of extra divergence ranging from 0 to 100% of the branch’s length is added (Fig. S2). At the end of each simulation, we reconstructed the simulated protein families and estimated their branch of origin with our pipeline for TRG detection (see above). Each simulated family that is not assigned to the branch root of the focal genus tree is a false positive simulated TRG family whose age has been underestimated because homologs are highly diverged. All real TRG families whose phylogenetic distances exceed the branch-specific threshold (under which a maximum of 5% false positives are expected) of the normal case scenario are excluded. Note that even compared to the worse case scenario, false positives cannot explain the total percentages of the observed TRGs (Fig. S3).

### Sequence properties

Codon usage and Codon Adaptation Index (CAI) values for protein coding sequences were calculated with the CAIJAVA program version 1.0 (Carbone et al. 2003) (which does not require any set of reference sequences) with 15 iterations. CAI for the intergenic sequences was calculated with codonW version 1.3 (http://codonw.sourceforge.net/) afterwards, based on codon usage of genes with CAI > 0.7 (previously estimated with CAIJava), so as to avoid any bias that may be present within intergenic regions.

The expected number of amino acids in a transmembrane region were calculated with the TMHMM program (Krogh et al. 2001). Disordered regions were defined as protein segments not in a globular domain and were predicted with IUPRED version 1.0 (Dosztányi et al. 2005).

Low complexity regions were detected with *segmasker* version 1.0.0 from the BLAST+ suite. Biosynthesis costs were calculated using the Akashi and Gojobori scores (Akashi and Gojobori 2002; Barton et al. 2010). GRAnd AVerage of Hydropathy (GRAVY) and aromaticity scores of each protein sequence were calculated with codonW version 1.3. Predictions of helices and sheets in protein sequences were obtained by PSIPRED version 3.5 (McGuffin et al. 2000) in single sequence mode. TANGO version 2.3 (Fernandez-Escamilla et al. 2004) was used to predict the mean aggregation propensity per residue for all proteins with the settings provided in the tutorial examples.

### Calculation of Coding Score

We built a binomial logistic regression classifier on a Coding class and a Non-coding class. The Coding sequences are genes conserved inside and outside of the focal genus. The Non-coding sequences corresponding to the +1 reading frame of intergenic regions in which in-frame stop codons were removed. All non-annotated regions were considered in the *Lachancea* genomes, while orthologous intergenic regions available at www.SaccharomycesSensuStricto.org where considered in the *Saccharomyces* genomes. Both classes have equal sizes (6000 sequences each), which are sampled to have approximately the same length distribution. The Coding Score (CS) is the model’s fitted probability for the Coding class. The classifier was trained on the following sequence feature data: frequencies of 61 codons, CAI, biosynthesis cost, percentage of residues in i) transmembrane regions ii) disordered regions ii) low complexity regions iv) helices v) beta sheets, hydrophobicity scores, aromaticity scores, mean aggregation propensity per residue and the GC.GC3 term:

GC.GC3 = abs(GC – GC3) / abs(GC - 0.5), where GC is the percentage of Guanine-Cytosine bases and GC3 is the percentage of Guanine-Cytosine bases at the 3^rd^ codon position.

Each feature value was normalized by subtracting the mean and dividing by the standard deviation. The binomial logistic regression classifier was constructed with the GLMNET R package version 2.0-2 (Friedman et al. 2010), with an optimized alpha value (0.3 and 0.4 for the *Lachancea* and for the *Saccharomyces*, respectively) estimated by testing on a separate validation set of coding and non-coding sequences, and keeping the value that minimized the class prediction error. The function *cv.glmnet* with the optimal alpha value was used on the training set to perform 10-fold cross-validation to select and fit the model that minimizes the class prediction error for a binomial distribution. Validation of the performance of the coding score is given in Fig. S4.

### Orientation analysis

Relative orientation of the 5’ transcribed element was considered for a given gene that was tagged either in opposing orientation (<– –>) if its 5’ neighbor is transcribed on the opposite strand or co-oriented (–> –>) if its 5’ neighbor is transcribed on the same strand. Only genes that do not overlap other elements on the opposite strand at their 5’ extremity (non-null intergenic spacer) were considered. Relative 5’ orientations were determined for *de novo* genes, conserved genes and tandemly duplicated genes. There are 925 and 580 tandemly duplicated genes in *Saccharomyces* and *Lachancea*, respectively are defined as paralogs that are contiguous on the chromosome. Among tandemly duplicated genes, 638 and 428 are co-oriented in *Saccharomyces* and *Lachancea*, respectively.

### Similarity searches in intergenic regions

For each chromosome, low complexity regions were first masked with *segmasker* version 1.0.0 and annotated regions were subsequently masked by *maskfeat* from the EMBOSS package version 6.4.0.0 (Rice et al. 2000). Similarity searches between all 6 frame translations of the masked chromosome sequences and the TRG protein sequences allowing for frameshifts were performed with the *fasty36 (Pearson et al. 1997)* binary from the FASTA suite of tools version 36.3.6 with the following parameters: BP62 scoring matrix, a penalty of 30 for frame-shifts and filtering of low complexity residues. Significant hits in at least two genomes (30% identity, 50% target coverage and an E-value lower than 10^−5^) within intergenic regions that are syntenic to a *de novo* gene were selected and their corresponding DNA regions were extracted. A multiple alignment was then performed and in-frame stop codons where searched in the phase whose translation is similar to the *de novo* gene product. All gaps that were not a multiple of three were considered as indels. In 16 cases, the enabling mutations from the ancestral non-coding sequence can be precisely traced forward based on the multiple alignment, as in Knowles and Mclysaght (Knowles and McLysaght 2009).

### Evolutionary analyses

For each TRG family with members in at least two different species, rates of synonymous substitutions (dS) and rates of non-synonymous substitutions (dN) were estimated from protein guided nucleotide alignments with the *codeml* program from the PAML package version 4.7 (Yang 2007). Pairwise analyses were done using the Yang and Nielsen model (Yang and Nielsen 2000). The relative rates dN/dS values were considered only if the standard error of dN and the standard error of dS were lower than dN/2 and dS/2 respectively and dS was lower than 1.5. Ancestral sequences were calculated with *baseml* from the PAML package version 4.7 using the REV model.

### Relative divergence estimates

Timetrees for both *Lachancea* and *Saccharomyces* were generated using the RelTime method (Tamura et al. 2012). For each genera, we selected 100 families of syntenic homologs present in every genome (in the 10 *Lachancea* or in the 5 *Saccharomyces*) for which the inferred tree has the same topology as the reference species tree (Scannell et al. 2011; Vakirlis et al. 2016). The concatenation of the protein-guided cDNA alignments of the family were given as input. As outgroup species, we used *S. cerevisiae* for the *Lachancea* and *Candida castellii* for the *Saccharomyces*. Divergence times for all branching points in the topology were calculated using the Maximum Likelihood method based on the Tamura-Nei model (Tamura and Nei 1993). 3^rd^ codon positions were considered. All positions containing gaps and missing data were eliminated. Evolutionary analyses were conducted in MEGA7 (Kumar et al. 2016).

### Recombination hotspots analysis

Recombination maps were retrieved from (Lam and Keeney 2015). The strains used to determine the recombination maps are those also used in this study (Scannell et al. 2011), so the same assembly has been used to map the Spo11 oligos for the recombination map and to detect *de novo* genes. This is not the case for *S. paradoxus*, because the recombination map is constructed for the YPS138 strain, which is quite divergent from the *S. paradoxus* strain CBS432 used to detect *de novo* genes, and for which only a low quality assembly is available.

### General procedures

All alignments were done with the MAFFT *linsi* executable (version 7.130b) (Katoh and Standley 2013). All statistical analyses were done in R version 3.1 (R Core Team 2014) with standard library functions unless otherwise noted. Phylogenetic distances from protein family alignments were calculated using *fprotdist* from the EMBOSS version 6.4.0.0 with the PAM matrix and uniform rate for all sites (- ncategories 1). The PAM matrix was chosen for consistency.

### Translation evidence

*De novo* genes in *S. cerevisiae* for which positive proteomic data are available are tagged as “with translation evidence”. This designation corresponds to protein products identified i) in MS-based proteome characterization studies, ii) as prey proteins in MS-based affinity capture studies, iii) in two-hybrid experiments, iv) as localized by fluorescent fusion protein constructs, v) as a substrate in phosphorylation assays, vi) identified in ribosome profiling experiments and/or vii) in protein-fragment complementation assays.

In *S. cerevisiae*, 13 out of the 302 CDS that we classified as spurious TRG show evidence of translation. Based on these *S. cerevisiae* data, the negative predictive value of the CS is 0.95 (13/302), *i.e.* there is a 95% probability that a spurious TRG, with a CS below our threshold is actually not a *de novo* gene.

### Mass spectrometry protocol

Single colonies of each species were inoculated in 3 mL YP + 2% Glucose and grown at 30**_**C. After 2 days growth, the liquid cultures were inoculated into 12mL of YP + 2% Glucose at 30**_**C and were grown until they reached an optical density of 1.0. Cultures were centrifuged at 4,000 RPM for 2 minutes and the supernatant was removed. The cells were washed in 1ml of 1M Sorbitol and centrifuged for 2 minutes at 15,000 RPM. The supernatant was removed and the cells were stored at −80 **°**C.

For each strain three biological replicates were analysed. Cells were resuspended in 100 μL 6 M GnHCl, followed by addition of 900 μL MeOH. Samples were centrifuged at 15,000 g for 5 min.

Supernatant was discarded and pellets were allowed to dry for ∼5 min. Pellets were resuspended in 200 μL 8 M urea, 100 mM Tris pH 8.0, 10 mM TCEP, and 40 mM chloroacetamide, then diluted to 1.5 M urea in 50 mM Tris pH 8(Carvunis et al. 2012). Trypsin was added at 50:1 ratio, and samples were incubated overnight at ambient temperature. Each sample was desalted over a PS-DVB solid phase extraction cartridge and dried down. Peptide mass was assayed with the peptide colorimetric assay (Thermo, Rockford).

For each analysis, 2 μg of peptides were loaded onto a 75 μm i.d. 30 cm long capillary with an imbedded electrospray emitter and packed with 1.7 μm C18 BEH stationary phase. Peptides were eluted with in increasing gradient of acetonitrile over 100 min (Hebert et al. 2014).

Eluting peptides were analysed with an Orbitrap Fusion Lumos. Survey scans were performed at R = 60,000 with wide isolation 300-1,350 mz. Data dependent top speed (2 seconds) MS/MS sampling of peptide precursors was enabled with dynamic exclusion set to 15 seconds on precursors with charge states 2 to 6. MS/MS sampling was performed with 1.6 Da quadrupole isolation, fragmentation by HCD with NCE of 30, analysis in the Orbitrap with R = 15,000, with a max inject time of 22 msec, and AGC target set to 2 x 10^5^.

Raw files were analysed using MaxQuant 1.5.2.8 (Cox and Mann 2008). Spectra were searched using the Andromeda search engine against a target decoy databases provided for each strain independently. Default parameters were used for all searches. Peptides were grouped into subsumable protein groups and filtered to 1% FDR, based on target decoy approach (Cox and Mann 2008). For each strain, the sequence coverage and spectral count (MS/MS count) was reported for each protein and each replicate, as well as the spectral count sum of all replicates.

The *de novo* genes that are translated are homogeneously distributed across the 10 *Lachancea* species (P=0.6, X^2^ test). The proportion of *de novo* genes detected (25/288, 8.7%) is significantly lower than that of conserved genes of similar length (66%), which by definition appeared before the most ancient *de novo* genes. This depletion could be due to *de novo* genes only being expressed under particular conditions or stresses that were not tested in our experiments. Conversely, MS/MS did not detect TRG eliminated as spurious by our procedure.

### Statistical Analysis

2-sided Wilcoxon rank-sum tests were performed to compare pairs of distributions of GC content and pairs of distributions of percentages of residues in disordered regions, at a P-value threshold of 0.05. Chi-square tests of association were used to compare gene orientations. Pearson’s correlation was used for the association of gene age - dN/dS and number of de novo origination events – substitutions per site

## Acknowledgments

This work was supported by the Agence Nationale de la Recherche (Grant GB-3G, ANR-10-BLAN-1606), the CNRS (Grant PEPS ExoMod Neo-Gene), the USDA National Institute of Food and Agriculture (Hatch Project 1003258), the National Science Foundation (Grant No. DEB-1442148), the DOE Great Lakes Bioenergy Research Center (DOE BER Office of Science DE-FC02-07ER64494) and the National Institutes of Health (Grant P41GM108538). CTH is a Pew Scholar in the Biomedical Sciences, supported by the Pew Charitable Trusts.

We thank M. Bailly-Bechet, A. Lambert, S. Lemaire, O. Namy, and C. Neuvéglise for help in the analysis of the results. We thank F. Barras, Y. Choquet, B. Dujon, R. Lavery, E. Rocha, and F.A. Wollman for fruitful discussions and critical reading of the manuscript.

*Data deposition:* Mass spectrometry Raw data is available on the chorus project (www.chorusproject.org) public experiment "*Lachancea de novo*" ID# 2884."

